# Considerations for the use of contrast agents with diffuse *in vivo* flow cytometry to detect circulating cancer cell populations

**DOI:** 10.1101/2025.07.28.667193

**Authors:** Joshua S. Pace, Grace Matheson, Gauri Malankar, Lei Wang, Melissa H. Wong, Summer L. Gibbs, Mark Niedre

**Affiliations:** Northeastern University, Department of Bioengineering, Boston, MA, USA; Oregon Health and Sciences University (OHSU), Department of Biomedical Engineering, Portland, OR, USA; OHSU, Knight Cancer Institute, Portland, OR, USA; OHSU, Department of Cell, Developmental and Cancer Biology, Portland, OR, USA

**Keywords:** *in vivo*-labeling, circulating tumor cell, fluorogenic contrast agents

## Abstract

**Significance:** Metastasis is a leading cause of cancer-related deaths. Disseminated circulating tumor cells (CTCs) through the bloodstream seed metastatic tumors at distant sites. Most methods for enumerating CTCs in humans clinically rely on drawing and analyzing small blood samples, but these may yield inaccurate estimates of CTC burden and cannot measure CTC changes over time. Identification and enumeration of CTCs for experimental or clinical purposes largely rely on marker-driven analyses by flow cytometry.

**Aim:** In principle, non-invasive fluorescence enumeration of CTCs directly in vivo could provide a more accurate method for enumerating CTCs. However, this will require specific contrast agent for CTCs. The goal of this work is to define characteristics of useful CTC contrast agents and perform preliminary testing of candidate contrast agents used for fluorescence guided surgery (FGS).

**Approach:** We evaluated a clinical small-molecule folate receptor targeted contrast agent (OTL38, pafolacianine), a fluorogenic pan-cathepsin contrast agent (VGT-309, abenacianine), and a set of custom designed, small-molecule prostate specific membrane antigen (PSMA) targeted contrast agents. We tested these contrast agents using *in vitro* cell culture models and in *in vivo* murine models.

**Results:** All tested contrast agents showed high uptake and labeling by target cell lines, but also small but significant labeling of non-cancer blood cells. Contrast agents that exhibited rapid clearance from circulation and the fluorogenic approach resulted in significantly reduced non-specific interfering background fluorescence signals.

**Conclusions:** Although the fluorescence contrast agents tested have properties useful for labeling of CTCs, as yet none exhibited the required high specificity. This resulted in some labeling of non-cancer blood cells which presented false-positive CTC counts. Improved contrast agent design and multiplexed use of more than one contrast agent may improve this specificity.

## 1 Introduction

Metastasis is spread of cancer from the primary tumor to distant organs or tissues and is the primary cause of cancer-related mortality (1, 2). One of the main pathways is hematogenous metastasis, wherein circulating tumor cells (CTCs) disseminate through the bloodstream. A small fraction may form secondary metastases (3–11). As such, CTCs are clinically significant and a focus of basic cancer research for over two decades (6, 7, 12, 13).

The gold standard for enumerating and characterizing CTCs is ‘liquid biopsy’, where a small volume of blood is collected from human subjects. CTCs can be isolated from whole blood by isolating peripheral blood mononuclear cells and capturing the cancer cells based on distinctive phenotypic traits such as epithelial cell surface protein expression, cell size, or mechanical characteristics (11, 14–17). Currently, CellSearch is the only FDA approved assay for enumerating and characterizing CTCs. Cells are identified as CTCs as those with positive epithelial cell adhesion molecule (EpCAM) and keratin while also being negative CD45 (18).

Despite significant promise, CTC liquid biopsy has yet to reach widespread clinical use, particularly as a predictive diagnostic method. Recent evidence suggests that this methodology can lead to inaccurate quantitative estimation of CTC numbers for several reasons. First, CTCs are rare and the drawn blood volume samples only a small percentage of total peripheral blood (PB) volume (e.g., 7.5 mL used for the CellSearch Clinical CTC specific assay compared to 5 L human PB volume) (19, 20). This small ratio may decrease quantitative accuracy and has a high likelihood of missing rare CTCs (i.e., low sensitivity), even if CTC numbers are stable and Poisson counting statistics are assumed (21, 22). Second, because of their short half-life in circulation, CTC numbers can fluctuate over short time periods (23). Moreover, CTCs levels exhibit diurnal or circadian patterns (24–26). Finally, the location of the blood draw relative to the tumor and major draining vessels can affect CTC counts (27). As a result, we and others have explored the use of non-invasive optical techniques for *in vivo* CTC enumeration have been explored (28–35).

We developed Diffuse *in vivo* Flow Cytometry (DiFC), an emerging fluorescence-based method for detecting and counting CTCs in murine models of cancer metastasis (36). Unlike microscopy-based approaches, DiFC uses a fiber-optic based optical probe to probe deeper tissue and large blood vessels using highly scattered light. Using a purpose-made optical instrument and signal processing algorithms, DiFC enables real-time detection and enumeration of the fluorescence signal from labeled CTCs. We have previously applied DiFC for longitudinal studies of CTC dissemination in mouse xenograft models using cancer cells expressing fluorescent proteins (21, 37, 38).

Although we originally envisioned DiFC as a small animal pre-clinical research tool, in recent years we have considered translation of DiFC to human cancer patients for fluorescence-based *in vivo* CTC enumeration methods (39). Specifically, using a series of computational, *in vitro*, and *in vivo* models we showed that DiFC could in principal support detection of a well-labeled CTC to tissue depths of up to 4 mm in a reflectance geometry in biological tissues (40, 41). In human anatomy, there are several potential large blood vessels such as the radial artery in the wrist that could be used to sample large amounts of circulating PB (>100 mL per minute) (42). In addition, this would require the use of red or near infrared (NIR) light, where light experiences lower attenuation and scattering compared to visible (blue-green) light.

However, a major challenge to clinical translation of DiFC is that CTCs would require direct fluorescent molecular labeling *in vivo*, unlike in murine models of metastasis where cancer cells can be labeled with fluorescent dyes or fluorescent proteins prior to injection. Our group and others have shown that specific populations of circulating cells can be labeled *in vivo* using receptor-targeted contrast agents in preclinical murine models (29, 37, 43, 44). Preliminary data also indicated that CTCs could be labeled with a cancer-specific fluorescence contrast agent in human patient blood samples (45).

More recently, we developed a NIR version of DiFC, in part for its compatibility with emerging fluorescent molecular contrast agents for use in fluorescence guided surgery (FGS) of cancer (46, 47). Our rationale was that FGS molecular contrast agents that target cancer with high specificity for primary tumors may likewise permit specific labeling of CTCs. Moreover, many of these agents have progressed along the regulatory pipeline and are under evaluation in clinical trials (47, 48). OTL38 (Cytalux, pafolacianine) is a folate-receptor alpha (FRα) targeted NIR small molecule contrast agent that was recently FDA-approved in the United States for lung and ovarian cancer (49–51). We recently demonstrated that OTL38 could be used to label CTCs directly *in vivo* in immunocompromised mice and subsequently detected with DiFC (52).

In this work, we explore considerations for the use of molecularly targeted contrast agents for labeling of CTCs and detection with DiFC. We performed proof-of-concept testing with three promising clinical and preclinical stage NIR and red contrast agents for CTCs: i) OTL38, ii) VGT- 309 (abenacianine), which is a pan-cathepsin (B, L, S, X) activatable NIR contrast that in phase 2 clinical trials for lung cancer and was recently given Fast Track FDA designation and, iii) purpose-made prostate specific membrane antigen (PSMA)-targeted fluorescence contrast agents for CTCs which are in pre-clinical development by our team. We note that while OTL38 and VGT-309 were not specifically designed for CTC enumeration, they have been shown to have high affinity for several cancer types and their progress through the regulatory pathway made them attractive candidates. We also discuss broader contrast agent considerations for specific and bright *in vivo* enumeration of CTCs (38, 53–55)

## 2 Materials and Methods

### 2.1 Overview of the CTC Contrast Agent Labeling Problem

Current cancer-specific contrast agent design and development has been focused on fluorescence molecular imaging of cancer for FGS, where the goal is specific labeling of the solid primary and metastatic tumors. However, labeling of CTCs for external optical detection with DiFC (or other fluorescence sensing methods) requires different contrast agent properties. As illustrated in **Figure 1**, an ideal CTC contrast agent for DiFC would have the following characteristics:

1. *High affinity for CTCs*: The fluorescence contrast agent must be directed against one or more molecular targets associated specifically with the cancer of interest, such as cell surface receptor or enzymatic over-expression. The approach requires sufficiently bright cell labeling for external detection with DiFC. In strategy 1, CTCs would be labeled while in circulation, although the circulation half-life is known to be relatively short for many cancers (**Fig. 1a**). In strategy 2, tumor and metastases would be labeled with the contrast agent and then labeled CTCs subsequently shedding into circulation (**Fig. 1b**).
2. *Low non-specific uptake by non-cancer cells in the blood*: In addition to non-specific background noise obscuring weakly-labeled cells, non-CTCs in the blood may take up the contrast agent presenting false-positive ‘peaks’ on DiFC. This may occur while non-CTCs are in circulation, or when non-CTCs reside elsewhere in the body and then traffic into the DiFC field of view. In a practical situation, it is known that non-CTCs outnumber CTCs by several orders of magnitude (i.e., CTCs frequently number on the order of 1-100 CTCs/mL, whereas white blood cells (WBC) number on the order of 10^6^ cells/mL). As such, specificity for cancer cells is critical for this this application.
3. *Low non-specific background signal and noise*: Administration of a fluorescent contrast agent (e.g., intravenously) results in elevated signal background. While many contrast agents have been shown to clear from circulation in several hours, their clearance from tissue can be on the order of days (53, 54, 56). Although tissue background fluorescence can generally be subtracted, the noise in the vascular background generally cannot. As such, higher non-specific background signal results in increased noise that can obscure more weakly-labeled cells. We previously showed in L1210A-bearing mice with OTL38 that this increased background signal resulted in loss of CTC counts by approximately 60% compared to baseline noise (52). In principal use of labeling strategy 1 (**Fig. 1a**) presents a higher background signal, whereas strategy 2 presents a lower background signal (**Fig. 1b**).

**Fig. 1.**
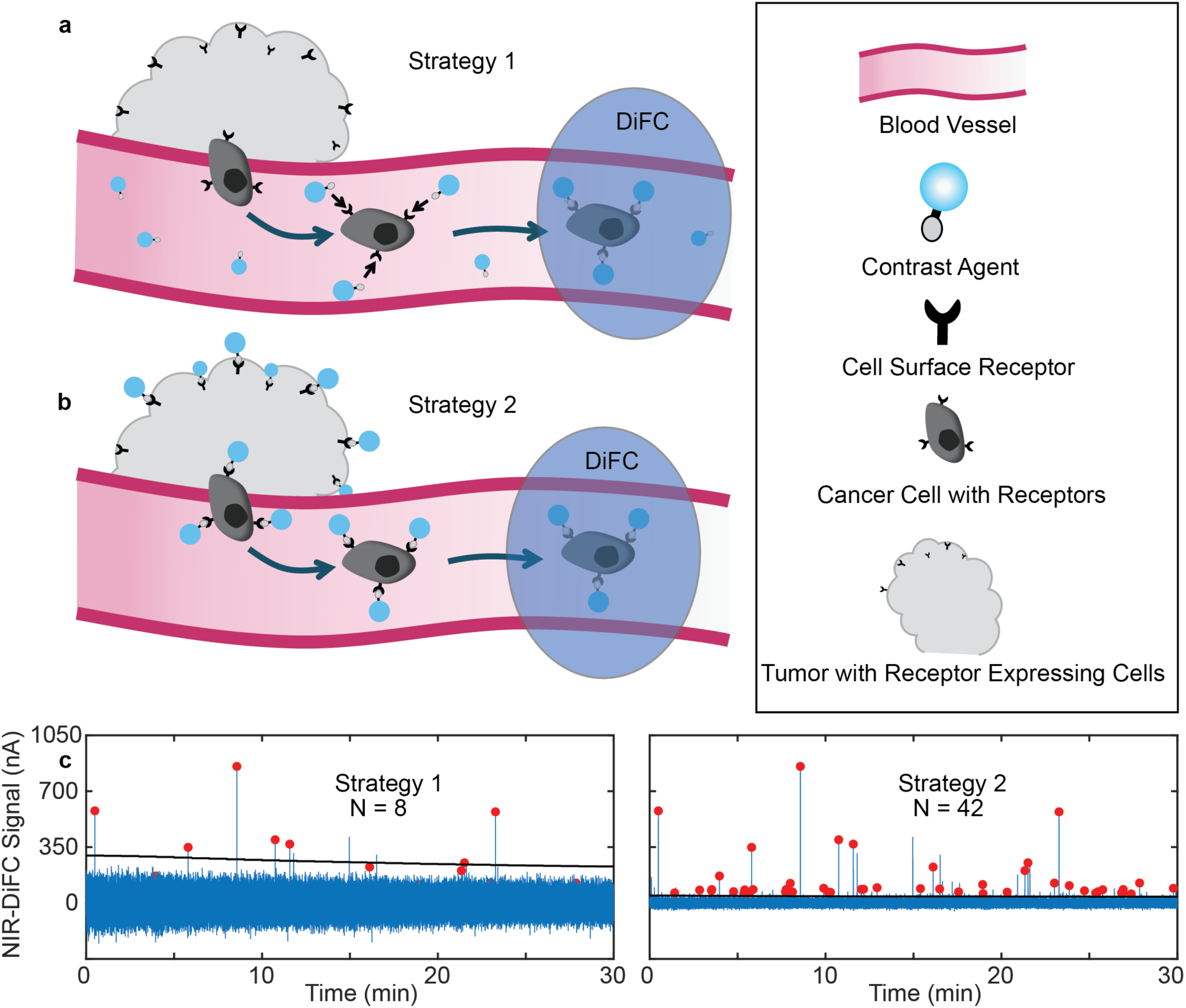
Labeling of CTCs at different timepoints after intravenous administration (IV) of contrast agent. **a** Closer to the time of contrast agent injection, cells in circulation can be labeled while there is free dye present (strategy 1). **b** Farther from the injection time, the contrast agent will be cleared from circulation and be accumulated in the tumor where cells shedding into the blood may still be labeled (strategy 2). **c** While cells can be labeled while in circulation, the presence of free contrast agent (and therefore an increased background) obscures dimmer labeled peaks (N = 8 detections). **d** After waiting a sufficient time for contrast agent clearance, the background will be closer to baseline and therefore dimmer labeled cells can be detected (N = 42 detections). Red circle labeled peaks are detected cells and the black horizontal line is the NIR-DiFC detection threshold.

### 2.2 Contrast Agents and Molecular Targets Tested

We tested three classes of contrast agents with differing molecular targets and regulatory approval stage as follows:

- *OTL38 – Folate Receptor Targeted*; OTL38 is a folate receptor alpha (FRα) -targeting fluorescent small-molecule (MW 1414.42 g/mol) contrast agent. FRα is frequently over- expressed in multiple types of cancer and has low expression in most normal tissues. OTL38 is a conjugate between a folate analog and S0456 dye (similar excitation and emission spectra to ICG) with a maximum excitation wavelength of 776 nm and stokes shift of 17 nm (57). OTL38 is also FDA approved for the use in fluorescence guided surgery of ovarian and lung cancer (49, 51, 58). We and others have shown that OTL38 (and earlier versions of the molecule) have high affinity for CTCs in blood, suggesting that it may be a promising CTC contrast agent. The OTL38 used in this study was generously provided by On Target Laboratories (West Lafayette, IN).
- *VGT-309 – Pan-Cathepsin Targeted*; VGT-309 (Vergent Bioscience, Minneapolis, MN) is a fluorogenic NIR (MW 2517.29 g/mol) contrast agent. VGT-309 uses ICG as the fluorophore with maximum excitation wavelength of 789 nm and stokes shift of 25 nm. When Cathepsin B, L, S, or X bind to the probe, the fluorophore is no longer quenched. Cathepsin activity is known to be higher in cancer than in surrounding normal tissue for certain organ sites (59). In addition to its affinity for cancer, we hypothesized that the fluorogenic properties of VGT-309 may present a very low background signal in the blood that may enhance CTC detection (see section 2.1). VGT-309 is also in phase II clinical trials for lung cancer (53, 54, 60). The VGT-309 used in this study was generously provided by Vergent Biosciences.
- *PSMA-0X – Prostate Specific Membrane Antigen Targeted*; We also tested several custom small-molecule fluorescent probes developed by our team that target the PSMA cell surface receptor. PSMA is known to be over-expressed in prostate cancer, but is also over- expressed in hepatocellular, glioblastoma, lung cancer and other tumor types. These probes used Cy5 (red) or Cy7 (NIR) fluorophores. The targeting molecules can be adjusted to alter binding and pharmacokinetics characteristic of the molecule. In this study, we used red and NIR versions of -02 and -04 variants, which were observed to have relatively slow and fast clearance kinetics, respectively.

### 2.3 Cell Lines Used

#### Folate Receptor Over-Expressing Cells

SK-OV-3 is an immortalized human ovarian cancer adherent cell line that naturally expresses FR (cAP-0054GFP; Angio-Proteomie, Boston, MA). IGROV-1 is an immortalized human ovarian cancer adherent cell line that naturally expresses FR (SCC203; Sigma-Aldrich, St. Louis, MO). All cell lines were cultured in RPMI 1640 folic acid deficient media (Gibco 27016021; ThermoFisher Scientific, Waltham, MA) supplemented with 10% fetal bovine serum (Gibco 16000044; ThermoFisher Scientific) and 1% penicillin/streptomycin (Gibco 15140122; ThermoFisher Scientific).

#### Cathepsin Over-Expressing Cells

Lewis Lung Carcinoma (LLC) that are known to have elevated Cathepsin activity and were also genetically modified to express green fluorescent protein (LL/2-Fluc-Neo/eGFP-Puro, Imanis Life Sciences, Rochester, MN) were utilized (61). Cells were cultured in DMEM (10-013-CV; Corning, Tewksbury, MA) with 10% fetal bovine serum (Gibco 16000044; ThermoFisher Scientific, Waltham, MA), 1% penicillin-streptomycin (Gibco 15140122; ThermoFisher Scientific), 2 μg/mL puromycin (Gibco A1113803; ThermoFisher Scientific), and 1.25 mg/mL G418 (Gibco 10131035; ThermoFisher). Addition of G418 antibiotic continuously selected for GFP-expressing LLC cells. 4T1 murine breast cancer cells (CRL-2539; ATCC) that express cathepsins were cultured in RPMI 1640 ATCC modified media (Gibco A1049101; ThermoFisher Scientific) supplemented with 10% fetal bovine serum (Gibco 16000044; ThermoFisher Scientific).

#### PSMA Over-Expressing Cells

LNCaP (CRL-1740; ATCC) and the derivative C4-2 (CRL-3314; ATCC) human prostate cancer cells that express PSMA were cultured in DMEM (10-013-CV; Corning) with 10% fetal bovine serum (Gibco 16000044; ThermoFisher Scientific). All cell lines were incubated at 37°C with 5% CO2. For all cancer cell experiments, cells were first collected from T-75 tissue culture treated flasks (FB012937; Thermofisher Scientific). For the adherent cell lines, the cell culture media was aspirated and then 6 mL TrypLE Express (Gibco 12604021; ThermoFisher Scientific) was added for 5 minutes to get the cells into suspension.

### 2.4 Measurement of Contrast Agent Specificity in Suspensions of Peripheral Blood Mononuclear Cells In Vitro

To study specificity of the different contrast agents for cancer cells in the presence of other types of blood cells that may bind or scavenge the contrast agents, we studied complex suspensions of human PBMCs (which include dendritic cells, monocytes, and lymphocytes). Peripheral Blood Mononuclear Cells [PBMCs, (PCS-800-011; ATCC)] were thawed and prepared according to ATCC instructions for *in vitro* non-specific contrast agent uptake studies. Mixtures of 1:1000 CFSE stained cancer cells to PBMCs 10^6^ PBMCs and 10^4^ CellTrace CFSE stained cancer cells were suspended in 1 mL of PBS. The contrast agent concentration, incubation time, and corresponding cancer cell line used is described in **Table 2**. All cell solutions were washed with fresh PBS twice and then analyzed by fluorescence Flow Cytometry (see **Section 2.8**). All samples were repeated at least N = 3 times with a minimum of 50,000 fluorescent count events collected.

**Table 1.** Contrast Agent type and stage of clinical approval.

| Contrast Agent | Probe Type | Target Molecule(s) | Regulatory Status |
| --- | --- | --- | --- |
| OTL38 | Small molecule | FRα | FDA approved |
| VGT-309 | Fluorogenic | Pan-Cathepsin | Phase II trial |
| PSMA-02,04 | Small molecule | PSMA | Pre-clinical |

**Table 2.** Concentration and incubation time used to pre-label cells with the fluorescent contrast agents used in this study.

| Contrast Agent | Cancer Cell Line | Concentration (nM) | Incubation Time (hr) |
| --- | --- | --- | --- |
| OTL38 | IGROV-1 | 200 | 1 |
| VGT-309 | LLC | 10000 | 2 |
| NIR-PSMA-02,04 | LNCaP | 250 | 1 |

### 2.5 Measurement of Contrast Agent Specificity in Mouse Blood Ex Vivo and in Mice In Vivo

Mice were handled in accordance with Northeastern University’s Institutional Animal Care and Use Committee (IACUC) policies on animal care. Animal experiments were carried out under Northeastern University IACUC protocol #24-0207R. All experiments and methods were performed with approval from and in accordance with relevant guidelines and regulations of northeastern University IACUC.

In DiFC measurements, elevated fluorescence background *in vivo* can obscure peaks from both true positive cancer cells and false positive cells. To study the non-specific labeling of cells circulating in the blood, we collected blood after IV contrast agent injection and sampled the blood with DiFC *ex vivo*. This allowed us to count non-cancer ‘false positive’ cells in the absence of interfering fluorescence background (noise) observed *in vivo* that would obscure DiFC detections. To investigate this, non-tumor bearing BALB/C mice (BALB/cAnNCrl; Charles River Labs, Cambridge, MA) were IV tail vein injection with an individual contrast agent, and blood was drawn via cardiac puncture at applicable timepoints, then flowed through a tissue mimicking flow phantom (**Fig. 2**), previously demonstrated (21, 36–38, 62). The phantom was made from high-density polyethylene (HDPE) block and has similar absorption and scattering properties to biological tissue (21). Microbore Tygon tubing (TGY1010C, Small Parts, Inc., Seatle, WA) was inserted into a hole drilled into the phantom at a depth of 0.75 mm. This depth was chosen to match the depth of the ventral caudal artery in the mouse tail (63).

**Fig. 2.**
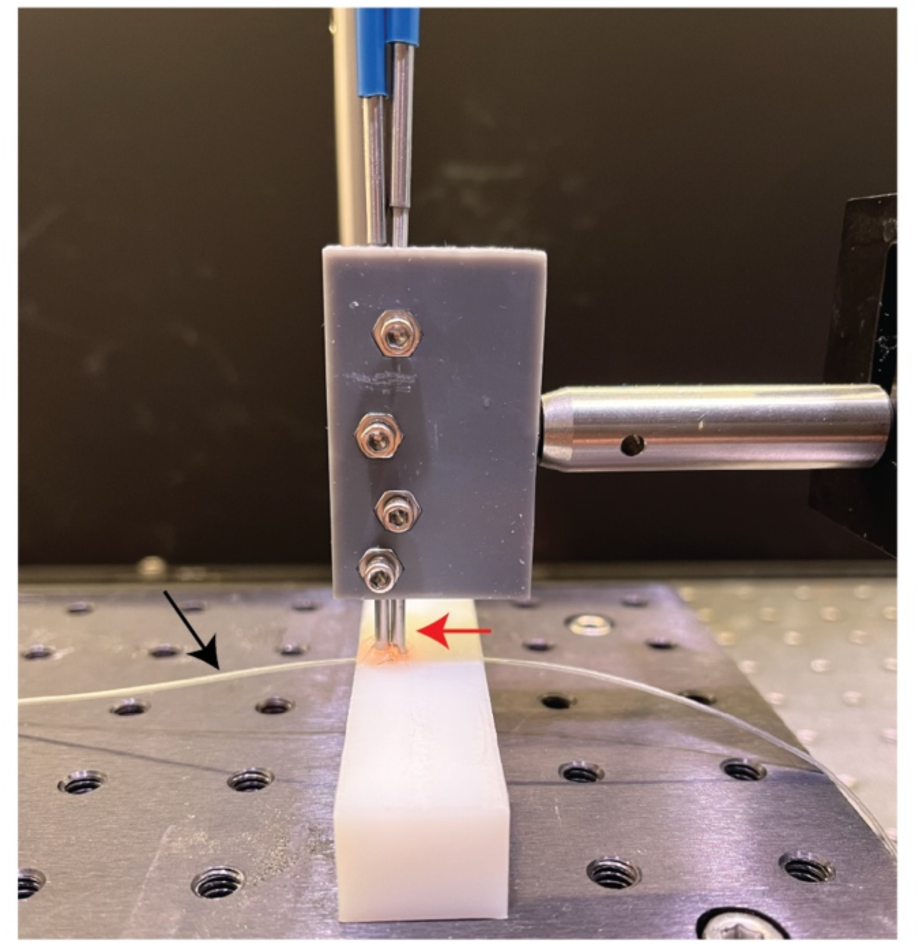
DiFC tissue mimicking phantom experimental setup. DiFC fiber probes (red arrow) are placed on top of a 0.75mm hole drilled through the optical phantom. Tygon tubing (black arrow) is fed through the hole and a syringe flows suspensions of cells through the phantom.

Concentration and time of DiFC scanning for each contrast agent are listed in **Table 3**. DiFC detections were counted and the number of non-cancer cells per mL of blood was calculated.

**Table 3.** Concentration and time after injection of DiFC scanning of the fluorescent contrast agents tested for non-specific cell uptake. Concentration and times are based on experiments we conducted or based on literature.

| Contrast Agent | Concentration (nmol) | DiFC scanning timepoints (hrs post injection) |
| --- | --- | --- |
| OTL38 | 1.8 | 3, 24 |
| VGT-309 | 10 | 24, 48 |
| Red-PSMA-02,04 | 10 | 4, 24 |

For *in vivo* experiments, OTL38 (1.8 nmol) and VGT-309 (10 nmol) were IV injected in non-tumor bearing athymic nude mice (NCR-nu/nu strain 553; Jackson Laboratory, Bar Harbor, ME) mice and DiFC was performed 24 hours post injection for 60 minutes on the mouse hindlimb.

### 2.6 Measurement of Contrast agent Background Signal In Vivo

DiFC collects highly scattered light from both the blood vessels and surrounding tissue. Mice were IV injected with a particular contrast agent and scanned with DiFC, with concentration and DiFC scan time for each contrast agent listed in **Table 4**. At each timepoint, DiFC measurements were collected for 1 hour and averaged.

**Table 4.** Concentration and time after contrast agent injection of DiFC scanning of the fluorescent contrast agents tested for fluorescence background measurements. Concentration and times are based on experiments we conducted or based on literature.

| Contrast Agent | Concentration (nmol) | DiFC scanning timepoints (hrs post injection) |
| --- | --- | --- |
| OTL38 | 10 | 3, 24 |
| VGT-309 | 10 | 2, 24 |
| Red-PSMA-02,04 | 10 | 4, 24 |

### 2.7 DiFC Instrumentation

#### 2.7.1 NIR-DiFC

The NIR diffuse *in vivo* flow cytometer (NIR-DiFC) used herein is the same instrument as previously introduced and characterized (46, 52). Briefly, the light source was a tunable pulsed laser (Mai Tai XF-1, Spectra Physics, Santa Clara, CA) with excitation wavelength set to 770 nm.

The light power at the sample was set to 25 mW. The collected light fibers were passed through an 810/10 nm bandpass emission filter (BP-em; FF01-810/10-25, IDEX Health and Science LLC) before being focused on to the surface of a photomultiplier tube (PMT; H10721-20, Hamamatsu, Bridgewater, New Jersey). Output signals from the PMTs were filtered with an electronic 100 Hz low pass filter, amplified with a low-noise current pre-amplifier (PA; SR570, Stanford Research Systems, Sunnyvale, CA), and then acquired with a data acquisition board (USB-6343 BNC; National Instruments, Austin, TX). NIR-DiFC uses custom-designed integrated fiber probe assemblies (EmVision LLC, Loxahatchee, FL). The design was an improvement compared to our previous fiber probe design (64) with better geometric collection efficiency and autofluorescence suppression. Each probe was constructed with 21 all silica low hydroxyl (OH) content 300 µm core 0.22 NA collection fibers, arranged in a 7-fiber inner ring and 14 fiber outer ring. The 21 collection fibers were arranged around a single fiber for laser delivery which is also an all silica 300 µm core low OH, 0.22 NA fiber.

A donut-shaped 807 nm long-pass filter (CLP; BLP01-785R, IDEX Health and Science LLC) was positioned in front of the 21 collection fibers to reject laser light and pass collected fluorescence light from the sample. The fibers, lens, and other optical components were placed inside a 2.4 mm outside diameter stainless steel needle tube.

#### 2.7.2 Red-NIR-DiFC Instrument

This work also utilized a newly developed Red-NIR DiFC to detect Cy5 and Cy7 fluorophores. The complete design was detailed but we note that only the Cy5 detection was use here. The instrument schematic is detailed in **Figure 3a**. A 638 nm laser (0638-06-01-0180-100; Cobolt, Solna, Sweden) with a 640/20 nm excitation filter (BP1-em; FF01-640/20-25, IDEX Health and Science LLC, Rochester, NY) was used for Cy5 excitation and the light power at the sample was

**Fig. 3.**
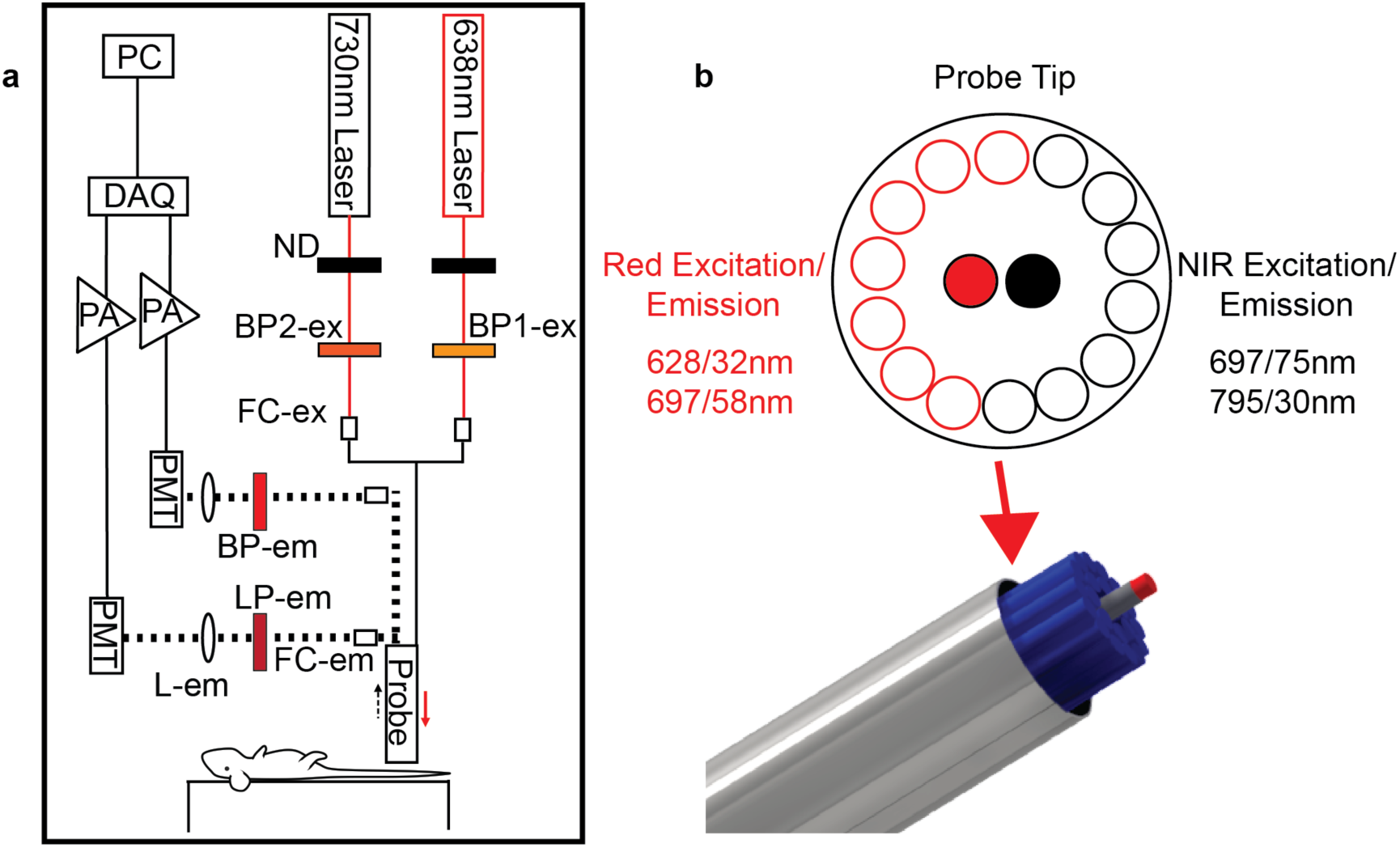
Red-NIR-DiFC Instrument and optical probe. **a** The instrument schematic and main components labeled (see text for details). **b** The optical fiber probe has two separate 300 µm excitation fibers surrounded by 14 total 300 µm collection fibers, 7 for Cy5 detection and 7 for Cy7 detection.

20 mW adjusted with a neutral density (ND) filter. A 730nm laser (0730-06-01-0050-100; Cobolt) with a 720/24nm excitation filter (BP1-em; FF01-720/24-12.5, IDEX Health and Science LLC) for Cy7 excitation and the light power at the sample was adjusted to 25 mW using a ND filter. Each laser was then separately coupled into source fibers with a collimation package (FC-ex; F240SMA-633, Thorlabs Inc., Newton, NJ). The output of the color separated probe collection fibers was collimated (FC-em; F240SMA-633, Thorlabs) and the light was passed through an 780 nm longpass emission filter for Cy7 detection (LP-em; ET780lp, Chroma, Bellows Falls, VT) and a 700/50 nm bandpass filter (BP-em; FF01-720/50-25, IDEX Health and Science LLC)before being focused on to the surface of a photomultiplier tube (PMT; H10722-20, Hamamatsu, Bridgewater, New Jersey) with a 30 mm focal length lens (L-em; 45363, Edmund Optics). The PMTs are powered by a power supply (C10709, Hamamatsu). Output signals from the PMTs were filtered with an electronic 100 Hz low pass filter, amplified with a low-noise voltage pre-amplifier (PA; SR560, Stanford Research Systems, Sunnyvale, CA), and then acquired with a data acquisition board (DAQ; USB-6343 BNC; National Instruments, Austin, TX).

Red-NIR-DiFC uses a custom-designed integrated fiber probe assembly as shown in **Fig. 3b** (EmVision LLC, Loxahatchee, FL). The probe was constructed with 14 all silica low hydroxyl (OH) content 300 µm core 0.22 NA collection fibers, 7 for Cy5 (668-726 nm collection filter) and 7 for Cy7 (780-810 nm collection filter). The 14 collection fibers are arranged around 2 fibers for 638 nm (with a 628/32 nm filter) and 730 nm (with a 697/75 nm) laser delivery which are also an all silica 300 µm core low OH, 0.22 NA fiber.

### 2.8 Flow Cytometry

Cell samples were analyzed using a Attune NXT flow cytometer (FC) (ThermoFisher Scientific). NIR fluorescence was collected using a 637 nm laser and a 780/60 nm emission filter. Red (Cy5) fluorescence was collected using a 637 nm laser and 670/14 nm emission filter. Green fluorescence was collected using a 488nm laser and a 530/30nm emission filter. Some cell samples were analyzed using a Cytoflex S FC (Beckman Coulter). NIR fluorescence was collected using a 638 nm laser and a 780/60 nm emission filter. Green fluorescence was collected using a 488nm laser and a 525/40nm emission filter. All samples were analyzed using FlowJo software and samples were gated for size and singlets based on corresponding cell populations.

## 3 Results

### 3.1 Contrast Agent Affinity and Specificity for CTCs in Suspensions of PBMCs In Vitro

Non-cancer cells in the peripheral blood outnumber rare CTCs by orders of magnitude (65). Hence, a major consideration for a usable CTC targeted contrast agent is high affinity and uptake of target CTCs but also low uptake for non-target leukocytes including peripheral blood mononuclear cells (PBMCs). To test this with OTL38, VGT-309, and PSMA-02 and -04 probes, we prepared suspensions of 1:1000 cancer cells-to-human PMBCs.

FR+ IGROV-1 cells were labeled with CFSE dye and then added to suspensions of PBMC cells. As shown in **Figure 4**, co-incubation with OTL38 resulted in labeling (increase in NIR fluorescence) for the target IGROV-1 cells but also resulted in some non-specific uptake by PBMC cells. In **Fig. 4b**, the vertical threshold represents ‘positive labeling’ by OTL38 measured with a benchtop FC instrument. Here, approximately 76.2 ± 11.5% of IGROV-1 cells were labeled by OTL38, as well as 3.3 ± 0.8% of PBMCs. The relative brightness for all cell lines tested in this experiment is summarized in **Fig. 4c**, along with our reference NIR fluorescent microsphere JGLI. The red vertical line represents the approximate *in vivo* DiFC detection threshold, which was previously determined relative to JGLI fluorescence intensity (46, 52).

**Fig. 4.**
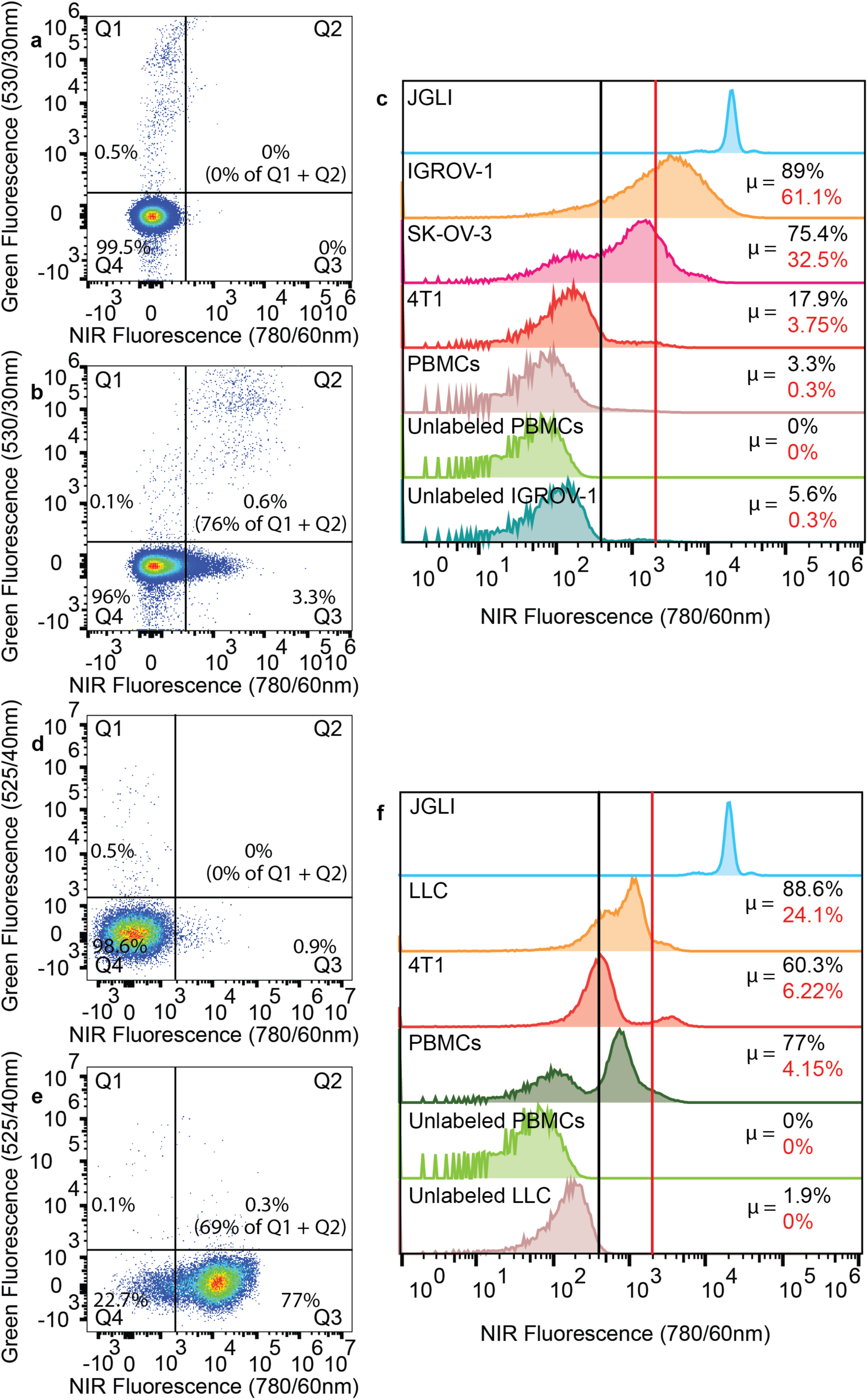
Flow Cytometry analysis of contrast agent labeled cancer cells and peripheral blood mononuclear cells (PBMCs). Suspensions of 1:1000 IGROV-1 ovarian cancer cells to PBMCs were incubated **a** without and **b** with OTL38. **c** Different FR expressing cancer cell lines and PBMCs incubated with OTL38 compared to unlabeled PBMCs, cells, and reference µspheres (JGLI). Labeled cell threshold (black line) and NIR-DiFC detection threshold (red line) are shown and percentages of cell populations above the respective threshold are indicated in the same color. Suspensions of 1:1000 LLC lung cancer to PBMCs were incubated **d** without and **e** with VGT-309. **f** Different cathepsin expressing cancer cell lines and PBMCs incubated with VGT-309 compared to unlabeled PBMCs, cells, and reference µspheres (JGLI). Labeled cell threshold (black line) and NIR-DiFC detection threshold (red line) are shown and percentages of cell populations above the respective threshold are indicated in the same color.

Briefly, the amplitude of detected JGLI microspheres with DiFC and flow cytometry were compared. From this, we estimate that under ideal labeling conditions, 61.1 ± 8%, 32.47 ± 18.8%, 3.8 ± 2.6% of IGROV-1, SK-OV-3, and 4T1 were sufficiently brightly labeled for detection with DiFC. Although only 0.3 ± 0.1% of PBMCs exceeded the estimated DiFC threshold, the relative abundance of PBMCs compared to CTCs was nevertheless problematic. For example, extrapolating this to an *in vivo* measurement with 100 IGROV-1 CTCs per mL, sampling 1 mL of blood would present approximately 61 true positive counts, but 2600 false positive counts. Hence, even though OTL38 shows high specificity for FR+ CTCs, the labeled PBMCs implied that this would present an unacceptably high false positive rate for DiFC. While the detection threshold could be increased to reduce false positive counts, this would of course correspondingly reduce the number of true positive detections.

Cathepsin+ LLC cells (also first stained with CFSE dye to differentiate from PBMCs) were co-incubated with PBMCs and VGT-309. As shown in **Fig. 4e**, 69.7 ± 13.5% of LLC cells (Q2) and 77 ± 23.4% of PBMCs were labeled above background (black vertical line) in the presence of VGT-309. The relative brightness for all cell lines are summarized in **Fig. 4f**. We estimate that under ideal labeling conditions, 24.1 ± 21.3%, 6.2 ± 4.4% of LLC and 4T1, respectively were sufficiently labeled for detection with DiFC. Due to the high abundance of cathepsins in PBMCs, 4.3 ± 0.4% were sufficiently labeled for DiFC detection. Again, extrapolating this to an *in vivo* measurement with 100 LLC CTCs per mL, sampling 1 mL of blood would present approximately 24 true positive counts, but 43,800 false positive counts. For this application of labeling and detecting CTCs in the blood, a cathepsin targeted probe is not feasible.

PSMA+ LNCaP cells (also first stained with CFSE dye to differentiate from PBMCs) were co-incubated with PBMCs and PSMA-02 and -04. As shown in **Fig. 5b**, 73.5 ± 10.4% of LNCaP cells (Q2) and 4.4 ± 1.9% of PBMCs were labeled above background (black vertical line) in the presence of PSMA-02. The relative brightness for all cell lines tested with PSMA-02 is summarized in **Fig. 5c**. We estimate that under ideal labeling conditions, 17.7 ± 0.4%, 29.1 ± 6.3% of LNCaP and C4-2 were sufficiently labeled for detection with DiFC. 0.2 ± 0.1% of PBMCs were sufficiently labeled for NIR-DiFC detection. The DiFC detection threshold presented was measured for our current instrument and was relatively higher than that for the NIR system. In principle the probe design could be better optimized for Cy7 in the future studies. Again, extrapolating this to an *in vivo* measurement with 100 C4-2 CTCs per mL, sampling 1 mL of blood would present approximately 29 true positive counts, but 2300 false positive counts.

**Fig. 5.**
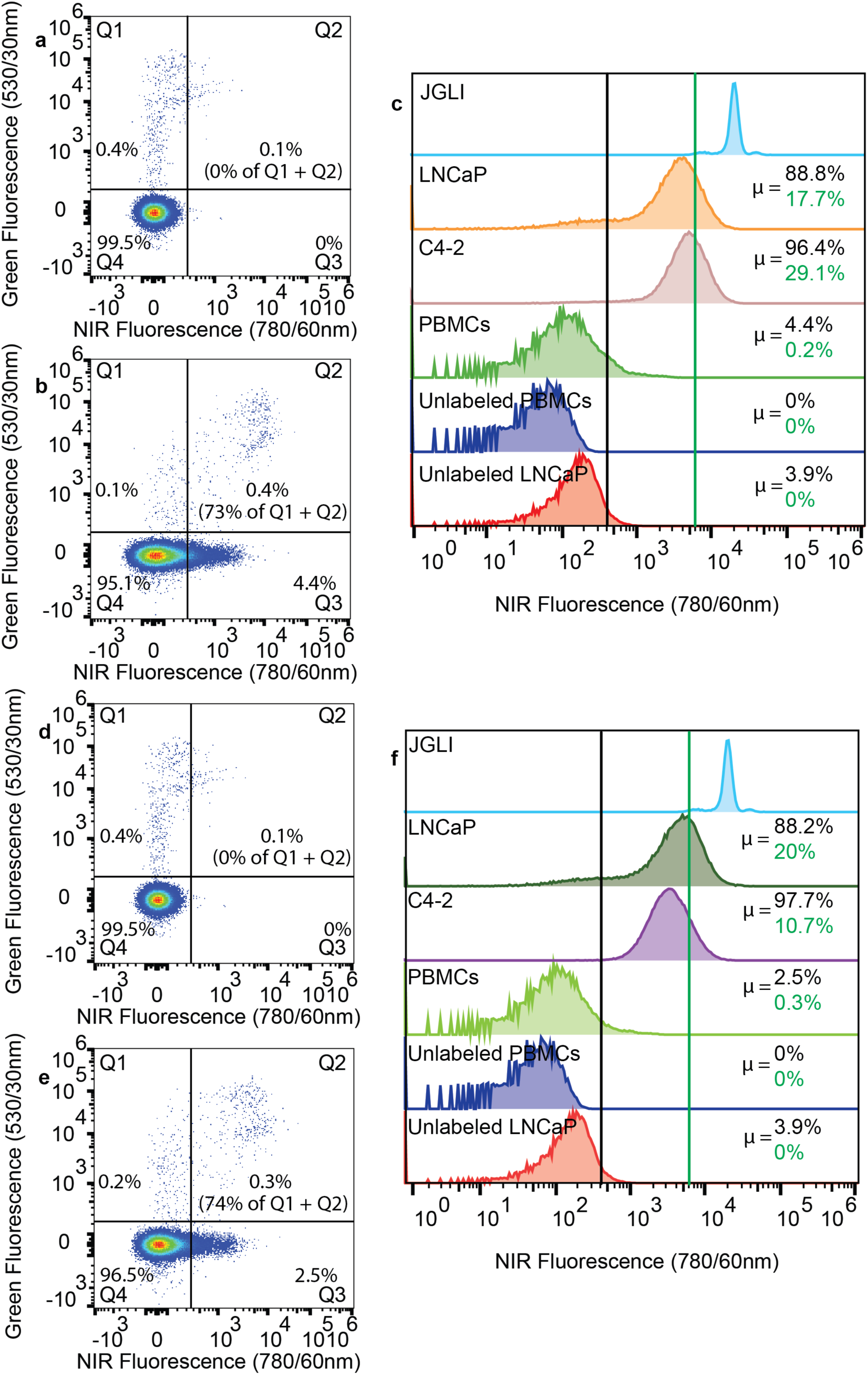
Flow Cytometry analysis of contrast agent labeled cancer cells and peripheral blood mononuclear cells (PBMCs). Suspensions of 1:1000 LNCaP prostate cancer cells to PBMCs were incubated **a** without and **b** with PSMA-02. **c** Different PSMA expressing cancer cell lines and PBMCs incubated with PSMA-02 compared to unlabeled PBMCs, cells, and reference µspheres (JGLI). Labeled cell threshold (black line) and Red-NIR-DiFC detection threshold (green line) are shown and percentages of cell populations above the respective threshold are indicated in the same color. Suspensions of 1:1000 LNCaP prostate cancer to PBMCs were incubated **d** without and **e** with PSMA-04. **f** Different PSMA expressing cancer cell lines and PBMCs incubated with PSMA-04 compared to unlabeled PBMCs, cells, and reference µspheres (JGLI). Labeled cell threshold (black line) and Red-NIR-DiFC detection threshold (green line) are shown and percentages of cell populations above the respective threshold are indicated in the same color.

As shown in **Fig. 5e**, 74.8 ± 8.2% of LNCaP cells (Q2) and 2.5 ± 0.7% of PBMCs were labeled above background (black vertical line) in the presence of PSMA-04. The relative brightness for cell lines tested is summarized in **Fig. 5f**. We estimate that under ideal labeling conditions, 19.7 ± 13.5%, 10.7 ± 8.6% of LNCaP and C4-2, respectively exceeded the NIR- DiFC detection threshold. 0.3 ± 0.1% of PBMCs exceeded the NIR-DiFC detection threshold. Again, extrapolating this to an *in vivo* measurement with 100 C4-2 CTCs per mL, sampling 1 mL of blood would present approximately 10.65 true positive counts, but 3200 false-positive counts.

### 3.2 Ex Vivo Contrast Agent Specificity in Mouse Blood

Although non-specific binding of contrast agents by PBMCs was observed, the *in vivo* labeling was more complex particularly considering labeling strategy 2 (**Figure 1**). As discussed in more detail in **Section 3.4** below, it is known that the contrast agent will clear from circulation after injection. Hence in addition to PBMCs, non-cancer cells anywhere in the body may in principle, take up the contrast agent and then happen to traffic to the bloodstream by the time the DiFC scan is performed presenting the possibility of a false-positive signal. On the other hand, it is conceivable that PBMCs that take up the contrast agent initially may clear from circulation before DiFC scanning.

To investigate this effect, non-tumor bearing immunocompetent BALB/C mice were IV injected with a contrast agent and collected blood was flowed through a phantom for *ex vivo* DiFC detection. This experimental approach was taken to remove the increased background fluorescence signal measured *in vivo* which could obscure detection of more weakly-labeled cells in the blood. In addition, there were additional sample preparation steps prior to passing the complete blood and heparin mixture through the optical phantom (as opposed to flow cytometry which generally requires depletion of erythrocytes).

In mice injected with OTL38, at both 3 and 24 hours we measured more than 600 cells/mL of blood detected (**Fig. 6a**). VGT-309 labeled over 1700 cells/mL of blood 48 hours post injection (**Fig. 6b**). Cathepsin activity in different types of immune cells could account for this high non-cancer cell uptake (66).

**Fig. 6.**
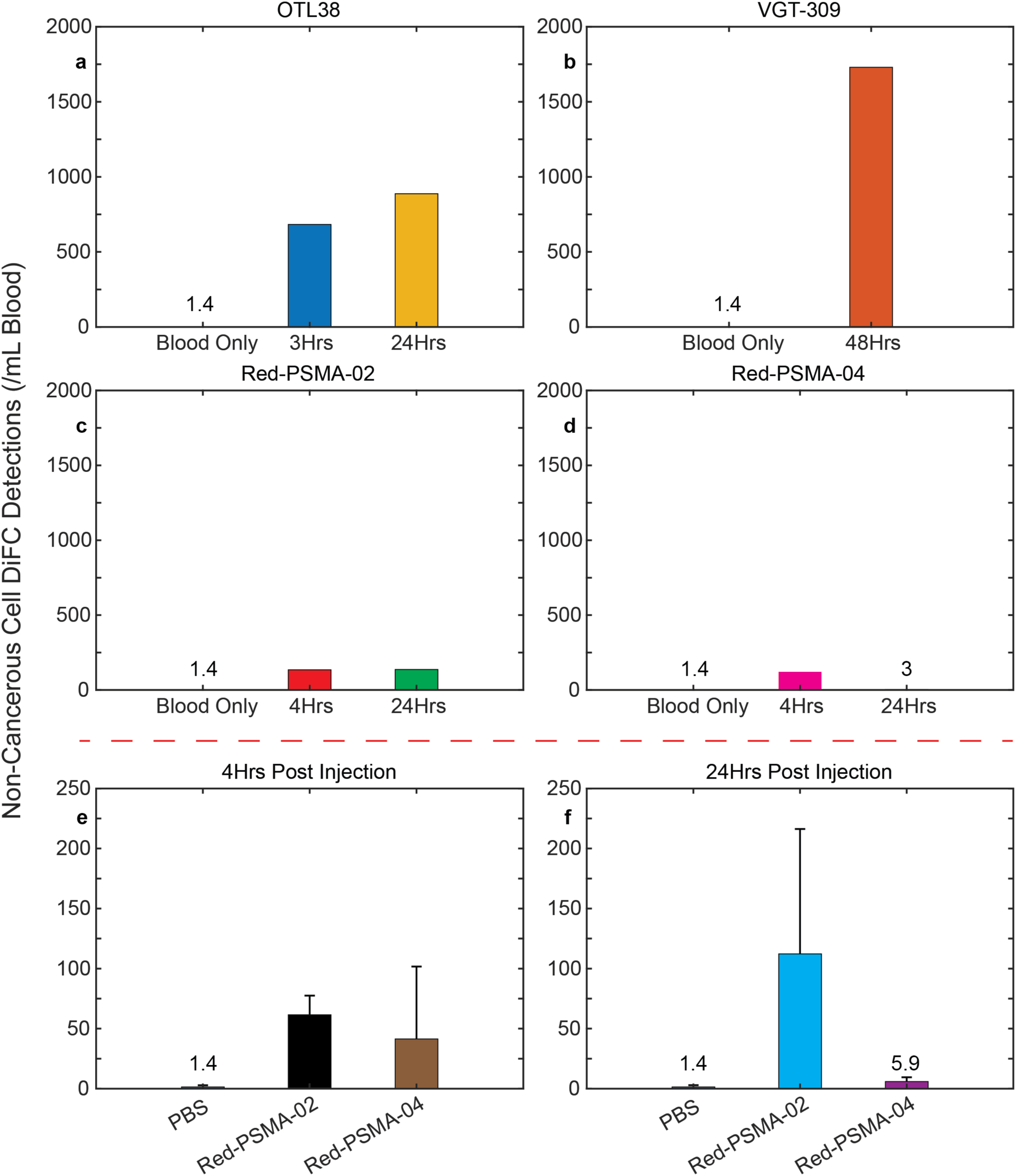
E*x vivo* DiFC detection of contrast agent labeled non-cancer cells in immune competent mice. BALB/C mice were IV injected with a contrast agent and timepoint specific collected blood was flowed through an optical phantom. **a** OTL38 had high immune cell false positive rates (683.1, 888.0 cells/mL blood) at 3 and 24 h post injection, which could be due to a more abundance of FR compared to other receptors found on cells. **b** VGT-309 was heavily taken up (1729.7 cells/mL blood) by non-cancer cells which could be due to the high abundance of cathepsins found in cells. **c** Red-PSMA-02 labeled 134.7 and 137.1 cells/mL blood at 4 and 24 h post injection respectively. **d** At 4 h post injection, Red-PSMA-04 labeled 121 cells/mL blood and there was only 3 cells/mL blood detected at 24 h. We note that error bars are not indicated because in most cases one measurement was performed to minimize the number of mice used. In an additional study of PSMA- 02 and -04 (N = 3, male CD-1 mice) **e** 4 h post injection still resulted in many false positives (61.5 ± 16 cells/mL blood for PSMA-02 and 41.4 ± 60.3 cells/mL for PSMA-04) **f** while at 24 h post injection PSMA-02 still labeled cells to a larger degree than PSMA-04 (112.2 ± 104 cells/mL blood and 5.9 ± 3.6 cells/mL blood respectfully).

With respect to our custom designed PSMA probes, we found that red-PSMA-02 resulted in labeling of more than 100 blood cells/mL at both 4 and 24 hours post injection (**Fig. 6c**). On the other hand, Red-PSMA-04 labeled 100 cells after 4 hours but only 3 cells 24 hours after injection (**Fig. 6d**). As we discuss in **Section 3.4** below, a lower uptake rate for PSMA-04 is consistent with our observation that the half-life of clearance from circulation for Red-PSMA-02 was significantly longer than that for Red-PSMA-04 (approximately 11 h for Red-PSMA-02 and 2.5 for Red-PSMA-04). Moreover, the lower number of non-target detections may be in part due to a more specific molecular target than either OTL38 or VGT-309. For example, it is known that immune and other cells exhibit a low expression of PSMA (67).

After initial promising results with the PSMA-02 and -04 probes, we selected those probes for further study with N = 3 repeats in male CD-1 immune competent mice. At 4 h post injection, PSMA-02 labeled 61.5 ± 16 cells/mL blood and PSMA-04 labeled 41.4 ± 60.3 cells/mL blood (**Fig. 6e**). 24 h post injection, PSMA-02 labeled 112.2 ± 104 cells/mL blood and PSMA-04 labeled 5.9 ± 3.6 cells/mL blood (**Fig. 6f**). These repeated results indicate that the faster clearing probe reduced the number of false positive detections at the longer DiFC scanning timepoint.

### 3.3 In Vivo Contrast Agent Specificity in Non-Tumor Bearing Control Mice

Although analysis of blood samples indicates that non-cancer cells were labeled with all the contrast agents we tested, this does not necessarily translate into detection *in vivo* with DiFC. Specifically, *in vivo* measurements present additional complications including attenuation of the fluorescence signal as it propagates through the biological tissue, as well as background fluorescence and noise. Hence, we performed DiFC on the hindleg of non-tumor bearing control mice, 24 hours after injection of OTL38 and VGT-309 as summarized in **Figure 7**. To briefly summarize our previously reported data analysis approach, 2 optical fiber probes were aligned approximately above the leg blood vessels of interest (**Fig. 7a**). Peaks were identified as labeled cells if there were sequential detections in the probes with the appropriate time delay due to the 3 mm separation between the probes. Peaks detected in only 1 probe were labeled as ‘unmatched’ and were not counted as a true cell.

**Fig. 7.**
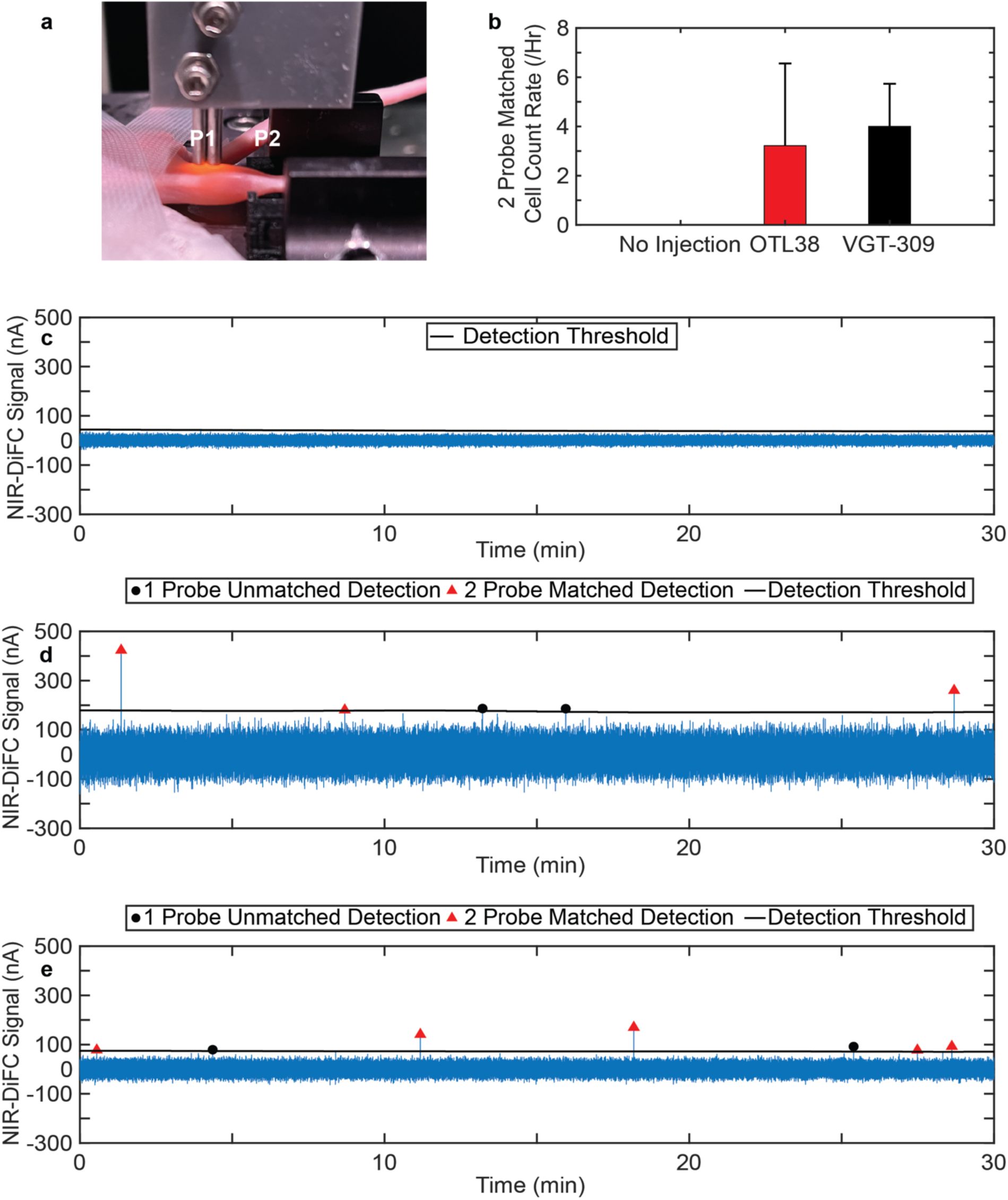
I*n vivo* detection of contrast agent labeled non-cancer cells with DiFC. **a** Two DiFC optical probes (P1 and P2) were placed on the skin over a blood vessel of interest which enabled ‘cell matching’ as a true cell (as opposed to motion) will be detected with P1 and P2 with a time delay due to the 3 mm spatial separation. **b** The count rate of detected non-cancerous blood cells (NCBCs) in non-tumor bearing mice injected with 10 nmol OTL38 (N = 3) was lower than 10 nmol VGT-309 (N = 3). Example DiFC data of non-tumor bearing mice **c** no cell detections were observed without contrast agent injection but were detected 24 h post IV injection of **d** 10 nmol OTL38 and **e** 10 nmol VGT-309.

As shown in **Fig. 7b**, the false positive detection rate was slightly (but not statistically) higher for mice injected with 10 nmol VGT-309 compared to 10 nmol OTL38 (an average false-positive rate of 4 ± 1.7 and 3.2 ± 3.4 per hour, respectively). By comparison, when DiFC was performed on control non-tumor bearing mice with no administered contrast agent, a lower background fluorescence was measured and no false positive matched detections for both OTL38 and VGT-309. Representative data is shown in **Fig. 7c**. 24 hours post 10 nmol OTL38 IV injection, scanning of non-tumor bearing mice resulted in true matched cell detections of non-cancerous cells (**Fig. 7d**). Similarly, 24 hours post 10 nmol VGT-309 IV injection, scanning of non-tumor bearing mice resulted in true matched cell detections of non-cancer blood cells (**Fig. 7e**) while the background signal was lower due to the fluorogenic property of VGT-309. This suggests that the combination of signal attenuation and noise contributions permitted detection of the most brightly-labeled non-cancer blood cells with DiFC *in vivo*.

We also note that this is a higher false-positive rate than we observed previously in nude mice with OTL38 (52). We attribute this difference to the longer time point (24 vs 3 h) as well as the higher concentration of OTL38 used in this study (10 vs 1.8 nmol).

### 3.4 In Vivo Measurement of Contrast Agent Background Signal

In addition to labeling CTCs, fluorescence from an injected contrast agent may create an elevated background signal in the blood and surrounding tissue which may obscure CTCs in DiFC measurements. To investigate how different contrast agents increase to the fluorescence background, DiFC *in vivo* measurements were collected (**Figure 8**). A DiFC optical fiber probe that delivers and collects light was aligned to a blood vessel (**Fig. 8a, picture 1**) then placed on the skin (**Fig. 8a, picture 2**). This resulted in highly scattered light being collected from both the blood vessel and surrounding tissue (**Fig. 8b**).

**Fig. 8.**
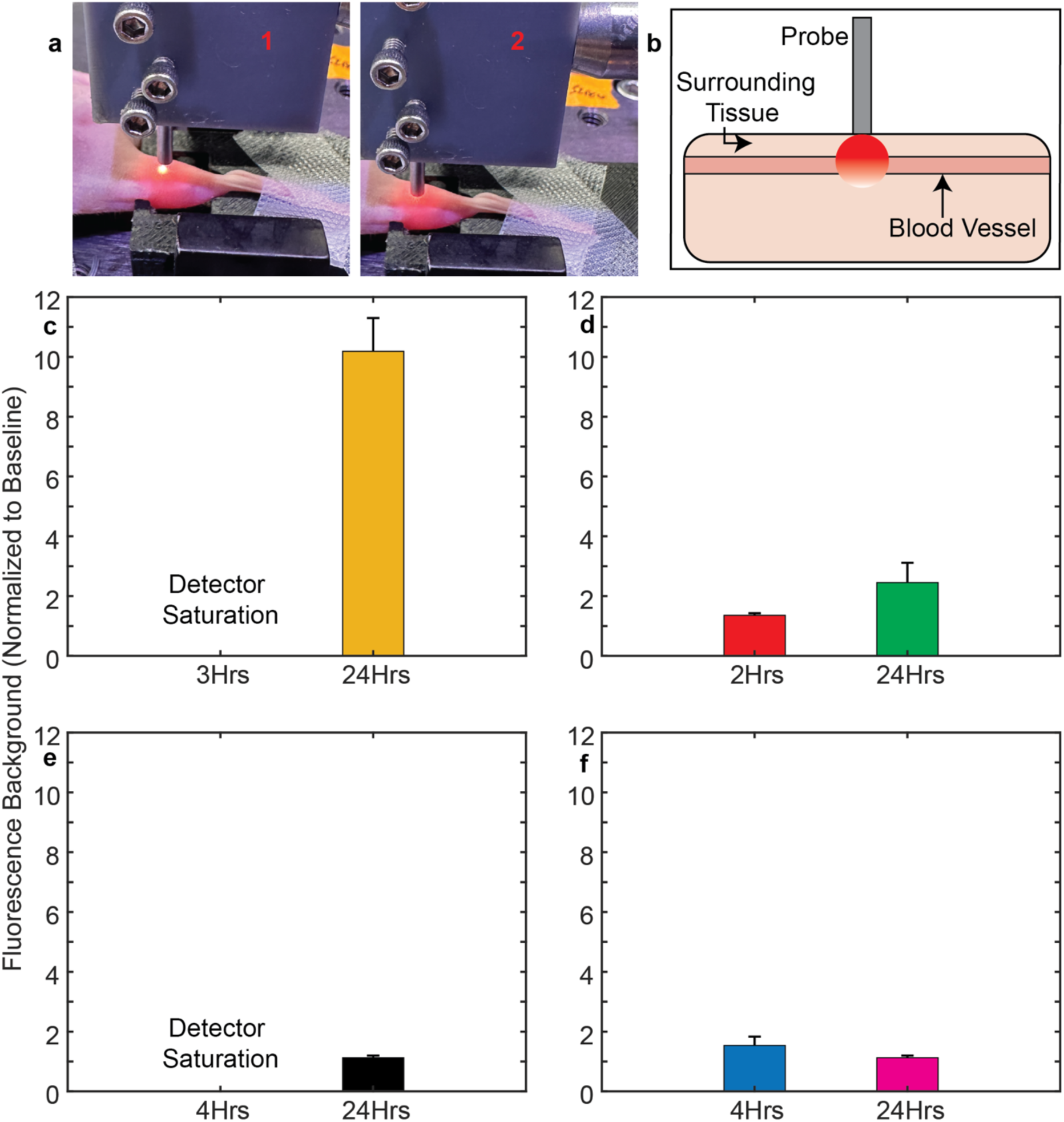
DiFC fluorescent background measurements after contrast injection in a mouse. **a** A DiFC optical probe that delivers and collects light (1) was placed on the skin directly above a blood vessel of interest (2). **b** Highly scattered light was collected from both the blood vessel and surrounding tissue. **c** After OTL38 injection, the fluorescence background increased and 3 h post injection the background was saturating the detectors (which makes the instrument unusable), 21 h later the background was approximately 10x baseline. **d** After VGT-309 injection, there was a small increase in background at 2 h and a larger increase at 24 h. This could be attributed to the fluorogenic property of VGT-309. **e** After red-PSMA-02 injection, the fluorescence background increases and 3 h post injection the background saturated the detectors (which makes the instrument unusable), 21 h later the background was back to baseline. **f** After injection of the faster clearing PSMA-04, the background was slightly above baseline at 4 h and was back to baseline at 24h.

*In vivo* measurements of were collected at specific timepoints after IV injection of contrast agent(s). 3 h post OTL38 injection, a measurement could not be obtained due to the instrument detectors saturating as a result of high NIR fluorescence from the contrast agent in the blood and surrounding hindlimb tissue (**Fig. 8c**). 21 hours later, the background signal was still approximately 10.2x baseline, suggesting that OTL38 may be retained in some tissues after clearance from plasma. These findings are similar to those reported for a human study in literature, where the plasma clearance half-life was determined to be in 2-3 h while in superficial skin, the half-life was 15 h with fluorescence still being detected 48 hours after injection (56).

2 h after VGT-309 administration, the background was approximately 1.4x baseline. 24 h post-injection, the background was higher 2.5x higher than at baseline (**Fig. 8d**). These lower background values may be attributed to the activable nature of VGT-309, where fluorescence is quenched until covalent binding of a cathepsin (53, 54, 60).

Injection of Red-PSMA-02, which clears from circulation in approximately 11 h, resulted in detector saturation at 4 h but back to baseline with only a 1.1x increase at 24 h. (**Fig. 8e**). Similarly with a faster clearing (approximately 2.5 h) version, Red-PSMA-04, background levels were 1.5x baseline 4 h post injection and were 1.1x at the 24 h.

## 4 Discussion and Conclusions

Although liquid biopsy and downstream analysis of CTCs has been shown to be a powerful tool, the methodology has seen limited adoption for clinical management of cancer patients. This is in part due to the small sample volume and infrequent analyses that results in CTC enumeration inaccuracy that comes from detection of rare cells. We and others posit that *in vivo* monitoring of CTCs provides a dynamic and continuous approach for non-invasive sampling of large circulating blood volumes (∼100 µL per minute in mice and 100 mL per minute in humans, respectively).

To begin developing this approach, development and validation of highly sensitive and specific fluorescent contrast agents for CTCs is essential. As we discussed, key properties for effective contrast agents are i) minimal fluorescence background, ii) high uptake by target cancer cells with iii) minimal uptake by non-cancer cells in the blood. Based on the studies herein, background fluorescence (i) was limited by both fast blood clearance of small-molecule probes, and use of fluorogenic (activatable) probes. Oral administration of contrast agents may further reduce this issue (compared to intravenous administration) since the probe would be more slowly released into the bloodstream. In addition, oral formulations of contrast agents would be more amenable for regular at-home or point of care screening for CTCs.

In terms of cell specificity, most contrast agents used in FGS target cell surface receptors that are overexpressed by cancer cells with lower (but non-zero) expression by other cell types. This principle generally works well for FGS since the measured signal is an aggregate of many different cells in the tumor and its microenvironment. Since CTCs are rare populations, false-positive uptake by non-cancer blood cells needs to be near zero for robust and accurate enumeration. For example, FR, PSMA, and cathepsins are expressed by immune cells and therefore can be falsely labeled and detected as we have shown here. On the other hand, because CTCs are already in the bloodstream issues with non-specific uptake (e.g., the enhanced permeability and retention effect) may be mitigated (68). We re-iterate that neither OTL38 or VGT-309 were developed specifically for labeling of CTCs, so that some uptake by non-cancer blood cells should not be surprising.

We also caution that although we attempted to optimize labeling conditions in each study, these results are specific to cell lines, mouse strains, contrast agents, concentrations, and timepoints tested. Moreover, results observed in mice may not directly translate to humans due to significant differences in contrast agent kinetics and CTC half-life. This said, these results suggest that a i) small molecule, ii) activatable, and iii) receptor targeted contrast agents are the most promising approach for robust labeling and noninvasive detection of CTCs. Specifically, low molecular weight probes appear to clear from circulation rapidly and exhibit reduced uptake by non-cancer blood cells. Fluorogenic (activatable) contrast agents are likely to reduce confounding fluorescent background signal.

Last, it is possible that a single CA may not be sufficiently specific for CTCs. We will also explore use of a second fluorescence tracer to improve CTC specificity, either to more clearly identify leukocytes (such as CD45) or as a non-specific paired agent (29, 69–71). Labeling and specificity results from this study are summarized in **Table 5**. For each category, a + indicates poor performance and a +++++ indicates excellent performance.

**Table 5.** Performance of contrast agents tested for uptake in cancer cells (*in vitro*), specificity to non-cancer cells (*ex vivo*), and the fluorescent background measured at timepoints after injection (*in vivo* and/or *ex vivo*).

|  | Cancer Cell<br>Labeling | Non-Target<br>Specificity | Fluorescent<br>Background |
| --- | --- | --- | --- |
| OTL38 | +++++ | ++ | + |
| VGT-309 | +++++ | + | +++++ |
| Red-PSMA-02 | +++++ | ++++ | +++++ |
| Red-PSMA-04 | +++++ | +++++ | +++++ |

Although more development of specific contrast agents and human-scale testing of DiFC is needed, robust and continuous in vivo monitoring of CTCs could have significant utility as a screening tool for management of cancer metastasis. For example, DiFC could allow early detection of early dissemination of CTCs during disease progression, detection of disease relapse, and monitoring of response to anti-cancer therapies. Beyond cancer, human-scale DiFC could have utility for cell enumeration in other applications such as CAR T-cell therapy, cell trafficking in injury response, or measurement of engineered circulating biosensors (72–74).

## Acknowledgements

This work was funded by the National Institutes of Health (R21CA246413) and a Kuni Foundation Discovery Grant (Gibbs/Wong/Niedre). The authors thank Prof. Philip S. Low and Dr. Madduri Srinivasarao of Purdue University, West Lafayette, IN, for the generous gift of OTL38 contrast agent used in this work. The authors thank Eric Benson and Vergent Bioscience for the generous gift of VGT-309 contrast agent used in this work.

## Code, Data, and Materials Availability

The datasets generated and analyzed during this work and the MATLAB code are available through the Pennsieve data sharing platform (DOI: 10.26275/btt6-puqw). Additionally, MATLAB processing code can be found in a Niedre Lab Github repository: https://github.com/mark-niedre/Contrast-Agents-and-DiFC-for-Detection-of-Circulating-Cancer-Cell-populations (DOI: 10.5281/zenodo.16371153).

## Author Bios

**Joshua Pace** received his BS degree in biomedical engineering from the University of Arizona in 2019. He received his PhD from the Northeastern University Department of Bioengineering in 2025.

**Summer Gibbs** received her PhD from Dartmouth College in Biomedical Engineering in 2008. She is a Professor of Biomedical Engineering at Oregon Health and Science University and a senior member of SPIE.

**Melissa Wong** received her PhD from Wake Forest University in Molecular Pathobiology in 1994. She is a Professor of Cell, Developmental and Cancer Biology at Oregon Health and Science University.

**Mark Niedre** received his PhD from the University of Toronto Department of Medical Physics in 2004. He is a Professor of Bioengineering at Northeastern University and a senior member of SPIE.

Biographies and photographs for the other authors are not available.

